# The structured demography of open populations in fluctuating environments

**DOI:** 10.1101/171884

**Authors:** Sebastian J. Schreiber, Jacob L. Moore

**Affiliations:** Department of Evolution and Ecology and Center for Population Biology University of California, Davis, USA

**Keywords:** Integral projection models, matrix models, open populations, stationary distributions, covariance formulas, sensitivity formulas

## Abstract

1. At the spatial scale relevant to many field studies and management policies, populations may experience more external recruitment than internal recruitment. These sources of recruitment, as well as local demography, are often subject to stochastic fluctuations in environmental conditions. Here, we introduce a class of stochastic models accounting for these complexities and provide analytic methods for understanding their long-term behavior.
2. The population state **n**(*x*) of these stochastic models is a function or vector keeping track of densities of individuals with continuous (e.g. size) or discrete (e.g. age) traits *x* taking values in a compact metric space. This state variable is updated by a stochastic affine equation **n**_*t*+1_ = **A**_*t*+1_**n**_*t*_ + **b**_*t*+1_ where is **A**_*t*+1_ is a time varying operator (e.g. an integral operator or a matrix) that updates the local demography, and **b**_*t*+1_ is a time varying function or vector representing external recruitment.
3. When the realized per-capita growth rate of the local demography is negative, we show that all initial conditions converge to the same time-varying trajectory. Furthermore, when **A**_1_, **A**_2_,… and **b**_1_, **b**_2_,… are stationary sequences, this limiting behavior is determined by a unique stationary distribution.
4. When the stationary sequences are periodic, uncorrelated, or a mixture of these two types of stationarity, we derive explicit formulas for the mean, within-year covariance, and auto-covariance of the stationary distribution. Sensitivity formulas for these statistical features are also given.
5. The analtyic methods are illustrated with applications to discrete size-structured models of space-limited coral populations, and continuously size-structured models of giant clam populations.

## Introduction

Recruitment corresponds to the addition of juveniles to a population through either local births or immigration events. When recruitment mainly occurs through local births, the population is closed, else it is open. At smaller spatial scales, which are often the scales at which empirical studies are conducted, open populations are common. This is especially common in species which only disperse during one life stage, such as plants with seed dispersal, insects with aerial dispersal, and marine organisms with larval dispersal [Hixon et al., 2002]. The simplest model of open populations are affine models, **n**_*t*+1_ = **An**_*t*_ + **b**, where **n**_*t*_ is the population state at time *t* (e.g. a vector of population densities), **A** describes the local demographic processes (e.g. a matrix accounting for survival, growth, reproduction, and emigration), and **b** describes external recruitment [Roughgarden et al., 1985, Pascual and Caswell, 1991, Caswell, 2008]. If this population relies on external recruitment to persist (i.e. the dominant eigenvalue λ of **A** is less than one), the population approaches a steady state **n̂** = –**A**^−1^**b** that depends on the interactive effects of external recruitment and local demography. This reliance on external recruitment may be due to a population living in a sink habitat where death rates exceed birth rates [Pulliam, 1988, Dias, 1996], or may be due to most newly born individuals emigrating. In either case, models of this form have been used successfully to understand the dynamics of open sessile populations with space limited recruitment [Roughgarden et al., 1985, Pascual and Caswell, 1991, Svensson et al., 2004], age-structured reef fish populations [Armsworth, 2002], and coral and clam populations with continuous size structure [Madin et al., 2012, Yau et al., 2014].

Open populations often experience environmentally driven fluctuations in external recruitment. If these fluctuations are serially uncorrelated and the population state is finite dimensional (i.e. **n**_*t*_ is a vector of densities), then one can models these populations by a first-order multivariate autoregressive, MAR(1), model: **n**_*t*+1_ = **An**_*t*_ + **b**_*t*_ [Reinsel, 2003, Ives et al., 2003, Gross and Edmunds, 2015, Cooper et al., 2015]. For these models, there is a unique stationary distribution provided that λ < 1, and explicit formulas for the mean and covariance matrix of this stationary distribution are well-known [Reinsel, 2003, Ives et al., 2003]. More recently, Gross and Edmunds [2015] developed sensitivity formulas for these means and covariances.

These MAR(1) models, however, do not account for fluctuations in local demography (i.e. **A** is time dependent), continuous population structure (e.g. size), or temporally correlated demographic fluctuations, though the importance of these population features is recognized. For example, Yau et al. [2014] used a continuous size-structured IPM to investigate how the degree of self-recruitment (used as a metric for the openness of the population) influenced management priorities of a giant clam fishery. Additionally, a series of matrix models was developed to account for space-limited recruitment in barnacle [Roughgarden et al., 1985, Hyder et al., 2001] and coral [Pascual and Caswell, 1991] populations. As a result of how space-limited recruitment was incorporated into the model structure, the amount of settlement influenced both **b** and **A**. These models were extended by Svensson et al. [2004, 2005] to investigate the relative importance of variability in recruitment, local survival, and migration. The models also accounted for correlated fluctuations in demography, through the use of randomly selected winter/summer periodic transition matrices. However, no analytic methods methods have been developed that account for these three biological features.

To address these fundamental biological complexities, we develop results for analyzing open population models accounting for (i) stationary and non-stationary fluctuations in local demography and recruitment, and (ii) any mixture of continuous and discrete population structure. Our results include conditions ensuring convergence of the processes for stationary and non-stationary environments, formulas for mean and covariance structure of the stationary distributions of the processes for any mixture of periodic and serially uncorrelated fluctuations, and sensitivity formulas for these statistical features. To highlight the broad applicability of our results, we apply them to discrete size-structured models of space-limited coral populations, and continuously size-structured models of giant clam populations.

## Models

Let 𝓢 be the set of individual states for the population of interest. For example, for a stage-structured matrix model 𝓢 = {1, 2, …, *k*} where states may correspond to a finite number of stages, ages, or spatial locations [Caswell, 2001]. Alternatively, for the simplest integral projection models (IPMs) [Easterling et al., 2000], 𝓢 = [*a*, *b*] corresponding to the continuum of sizes of individuals from the smallest of size *a* to the largest of size *b*. More generally, 𝓢 may correspond to a mixture of continuous and discrete structure. For example, for a size and age-structured population, 𝓢 = [*a*, *b*] × {1, …, *k*} where *k* is the maximal age of an individual [Ellner and Rees, 2006]. A common feature of all these examples is that 𝓢 is a compact metric space which we assume is the case for all the models discussed here.

To keep track of the densities of individuals, we have functions **n**: 𝓢 → [0, ∞) where **n**(*x*) is the “density” of individuals in state *x*. In general, we assume the functions n are continuous functions, but more general classes of functions are allowed as discussed the Appendices. Let **n**_*t*_(*x*) denote the density of state *x* at time *t*. To project the density function **n**_*t*_ forward in time, we have contributions due to individuals in the population, and contributions from outside of the population. To describe the endogenous contributions, let **A**_*t*+1_ be a linear operator taking nonnegative, continuous functions to non-negative, continuous functions. To describe the exogenous contributions, let **b**_*t*+1_(*x*) be the density of individuals in state *x* entering the population externally at time *t* + 1. Then the population model becomes

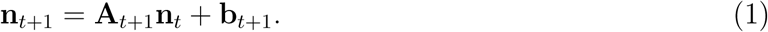

For a matrix model, **A**_*t*+1_ is a *k* × *k* non-negative matrix, **b**_*t*+1_ is a non-negative vector, and **A**_*t*+1_**n**_*t*_ + **b**_*t*+1_ corresponds to the usual matrix multiplication and vector addition. For the standard size-structured IPM, **A**_*t*+1_ is an integral operator of the form

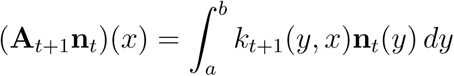

where *k*_*t*+1_(*y*, *x*) is a non-negative kernel describing the contribution of individuals of size *y* to individuals of size *x* at time *t* + 1, and **b**_*t*+1_ is a continuous function from [*a*, *b*] to [0, ∞).

Given a population density function **n**, let ‖**n**‖ be a norm corresponding to the total size of the population. For a matrix model, this norm is ‖**n**‖ = Σ_*s*_ **n**(*x*). For the standard IPM, two choices of this norm are the *L*^1^ norm ‖**n**‖ = 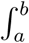 **n**(*x*)*dx* or the sup norm ‖**n**‖ = sup_*x*∈𝓢_ *n*(*x*). These norms on **n** also induces an operator norm on the operators **A**_*t*_ given by

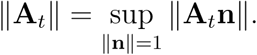

This operator norm corresponds to the largest change in the population size from time *t* – 1 to *t* that can be achieved by redistributing individuals across different states.

## Results: General Theory

We introduce three types of results. First, we present results for when the model converges to a time varying solution that is independent of initial conditions. Our condition for convergence applies to non-stationary as well as stationary environments. For stationary environments, this convergence is to a unique stationary distribution. Second, to understand the nature of these stationary distributions, we examine three special cases: periodic environments, temporally uncorrelated environments, and uncorrelated fluctuations around a periodic environment. Finally, we derive sensitivity formulas for properties of the asymptotic behavior in the three special cases.

### Convergence

Under mild assumptions, we introduce a natural condition that ensures the solutions of (1) converge to a time varying solution which is independent of the initial state of the system. The proof for this assertion follows from an argument of Brandt [1986] who studied a one-dimensional version of (1) in uncorrelated environments. Iterating the model (1) forward from time 0 to time *t* yields the solution:

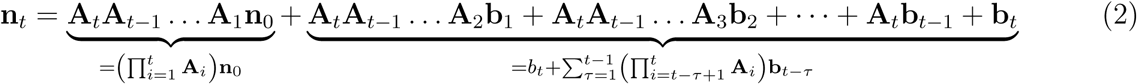

Importantly, only the first term depends on the initial state **n**_0_ of the population.

When this first term in (2) vanishes exponentially fast, we might expect that **n**_*t*_ converges to a time varying solution which is independent of the initial condition. To make this statement precise, define the *realized per*-*capita growth rate* of the population as

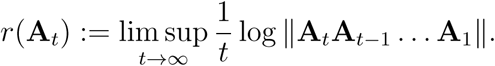

This term measures the maximal long-term per-capita growth rate of the population in the absence of immigration i.e. **b**_*t*_ = 0 for all *t*. If *r*(**A**_*t*_) < 0, then the population tends exponentially toward extinction in the absence of immigration. Namely, the population is a *sink population*. If *r*(**A**_*t*_) > 0, then the population asymptotically increases at an exponential rate *r*(**A**_*t*_) and is a *source population*.

For source populations, key properties of long-term dynamics, such as the long-term population growth rate, stable state distribution, and reproductive values, are determined by the “immigration-free” dynamics **n**_*t*+1_ = **A**_*t*+1_**n**_*t*_ for which classical results in stochastic demography apply [Tuljapurkar, 1990]. Hence, our results focus on the case of sink populations and we assume that *r*(**A**_*t*_) < 0. Furthermore, we assume that sup_*t*_ ‖**b**_*t*_‖ < ∞. Under these assumptions, we show in Appendix S1 that all population trajectories converge exponentially fast toward one another. That is, given two population trajectories **n**_*t*_ and **ñ**_*t*_ corresponding to initial conditions **n**_0_ and **ñ**_0_,

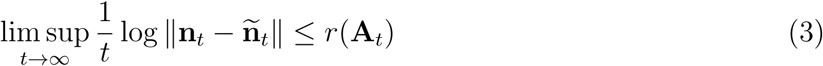

In particular, all population trajectories asymptotically approach the trajectory corresponding to the zero initial condition **n**_0_ = 0:

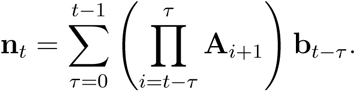

From a computational point of view, this result implies that it suffices to run the model once for any initial condition to understand its long-term behavior.

### Stationary environments

Assume that **A**_1_, **A**_2_, **A**_3_, … are a stationary and ergodic sequence of non-negative operators, and 𝔼[max{log ‖**A**_*t*_‖, 0}] < ∞. Kingman [1973]’s subadditive ergodic theorem implies there exists an *r* (possibly –∞) such that

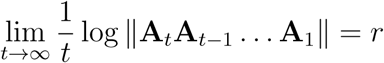

with probability one. In particular, the realized per-capita growth rate *r*(**A**_*t*_) equals *r* with probability one. This realized per-capita growth rate r corresponds to what Tuljapurkar [1990] calls the stochastic growth rate log λ_*S*_ of the population in the absence of immigration. In the mathematical literature, this common *r* value is known as the dominant Lyapunov exponent of the random sequence of operators [Ruelle, 1982].

If *r* < 0, then equation 3 implies that with probability one trajectories experiencing the same sequences of environmental conditions **A**_*t*_ and **b**_*t*_ but possibly different initial conditions converge to one another exponentially fast. Furthermore, from the ensemble perspective, we show in Appendix S2 that **n**_*t*_ converges exponentially fast to a unique stationary distribution provided that 𝔼[log ‖**b**_*t*_‖] < ∞. To define this stationary distribution, a standard probabilistic construction allows one to uniquely extend **A**_*t*_ and **b**_*t*_ to the past (i.e. …, **A**_–2_, **A**_–1_, **A**_0_, **A**_1_, **A**_1_,… and …, **b**_-2_, **b**_-1_, **b**_0_, **b**_1_, **b**_2_, …) such that they are stationary and ergodic. Then the unique stationary distribution is given by

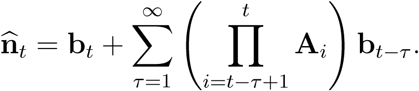

In the case of environments that are uncorrelated (i.e. independent and identically distributed), periodic, or a mixture of these types, we can say more about the stationary distribution.

#### Uncorrelated environments

Assume that **A**_1_, **A**_2_, … is an independent and identically distributed sequence of non-negative operators, and **b**_1_, **b**_2_, … is an independent and identically distributed sequence of density functions. In this case, we can write down an explicit, easily computed expression for the first- and second-order moments of the stationary distribution. Define λ̄ to be the dominant eigenvalue of Ā: = 𝔼[**A**_*t*_] and *r̄* = log λ̄. If *r̄* < 0 and 𝔼[‖**b**_*t*_‖] < ∞, then

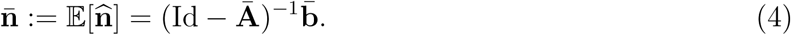

where **b̄** = 𝔼[**b**_*t*_] and Id denotes the identity operator: Id **n** = **n** for all **n**. As *r̄* > *r*, the existence of the stationary distribution isn’t sufficient to ensure that its mean is well-defined. In fact, when *r̄* > 0 > *r*, 𝔼[‖**n̂**‖] is infinite. We illustrate this phenomena with a simple scalar model in the examples section.

The covariance of **n̂**, denoted by **Cov**[**n̂**], is given by 𝔼[**n̂** ⊗ **n̂**] where ⊗ corresponds to the tensor product. In Appendix S3, we show that the covariance is well-defined when the spectral radius of 𝔼[**A**_1_ ⊗ **A**_1_] is less than one. When this spectral radius is less than one, 𝔼[**A**_1_ ⊗ **A**_1_] is invertible and

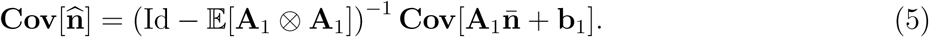

In the finite dimensional case, we can compute this covariance using the vec operation and the Kronecker product ⊗_*K*_ which corresponds to the tensor product in finite dimensions. Recall, for a *n* × *n* matrix *A*, the vec operation vec(A) is a column vector of length *n*^2^ given by concatenating the columns of *A*. For a *n* × *n* matrix *A* and *m* × *m* matrix *B*, the Kronecker product *A* ⊗_*K*_ *B* is the *nm* × *nm* block matrix given by

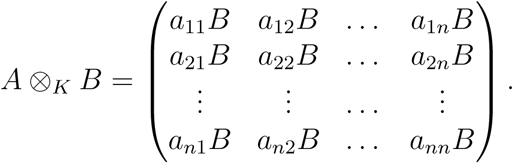

In terms of these matrix operations, the covariance matrix is given by (Appendix S3)

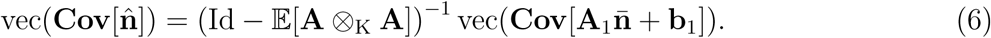

In the infinite dimensional case, one can apply (6) to the finite-dimensional matrix approximations of **A**_*t*_ and **b**_*t*_.

Even if **A**_*t*_ and **b**_*t*_ are uncorrelated in time, the population densities **n**_*t*_ typically will exhibit temporal correlations whenever there are overlapping generations. To characterize these temporal correlations, the covariance between **n̂**_*t*_ and **n̂**_*t*+*τ*_ equals

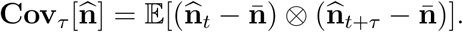

For *τ* ≥ 1, stationarity and independence imply

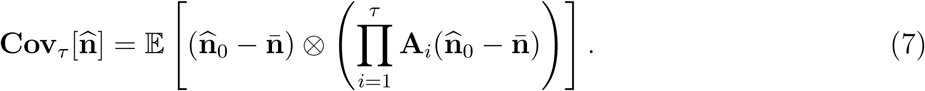

In finite dimensions, we can compute this auto-covariance with the following equation:

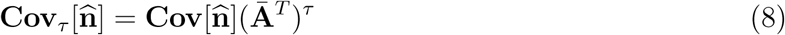

where ^*T*^ denotes the matrix transpose. As **Cov**[n̂] is only well defined when *r̄* < 0, (8) implies that the covariance terms, when they exist, decay exponentially fast with the length of the time lag *τ*.

Now assume that **A**_*t*_ and **b**_*t*_ depend on a parameter *θ*. The sensitivity of the mean **n̄** of the stationary distribution **n̂**_*t*_ is given by

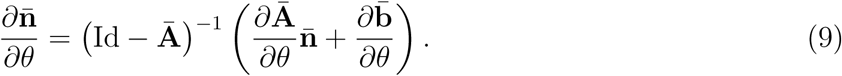

This sensitivity formula is the same as what Caswell [2008] found for deterministic matrix models of the form **n̄**_*t*+1_ = **Ān̄**_*t*_ + **b4**. The sensitivity of **Cov**[n̂] to *θ* equals

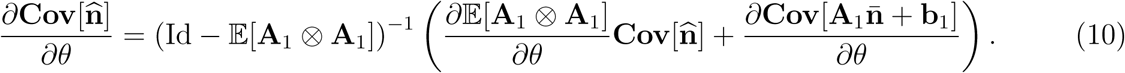

In finite dimensions, we can compute these sensitivities with the vec and Kronecker product:

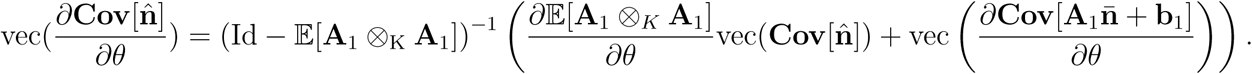

This finite dimensional sensitivity formulas agrees with what Gross and Edmunds [2015] found for finite-dimensional MAR models where there is no variation in the **A**_*t*_ matrices i.e. **A**_*t*_ = Ā for all *t*. In this special case, 𝔼[**A**_1_ ⊗_*K*_ **A**_1_] = **Ā** ⊗_*K*_ **Ā**. Finally, the sensitivity of the autocovariance for the finite dimensional models equals

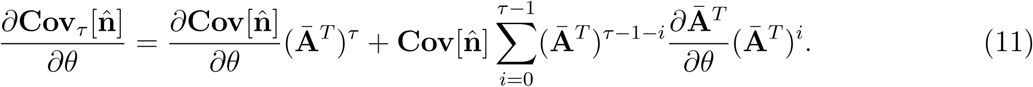

For the infinite dimensional models, we apply this formula to their finite-dimensional discretization.

#### Periodic environments

Consider a periodic environment with period *T*. Then **A**_*t*+*T*_ = **A**_*t*_ for all *t*. In the periodic environment *r*(**A**_*t*_) equals 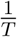 log λ where λ is the dominant eigenvalue of the period *T* operator **A**_*T*_ **A**_*T*–1_…**A**_2_**A**_1_. In particular, the condition *r*(**A**_*t*_) < 0 corresponds to the dominant eigenvalue being less than one. When this occurs, **n**_*t*_ converges exponentially quickly to the periodic trajectory (Appendix S4) given by

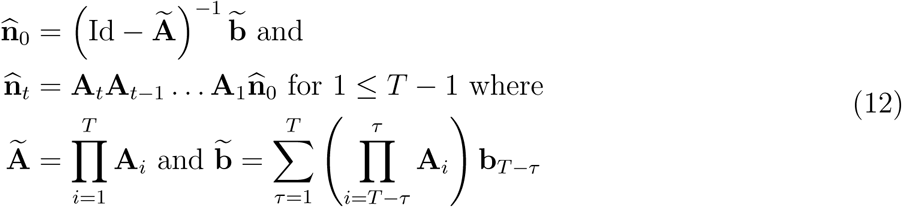

Now let *θ* be a parameter in the model. The sensitivity of the periodic solution, **n̂**_0_, **n̂**_1_, …, **n̂**_*T*–1_, to *θ* is given by

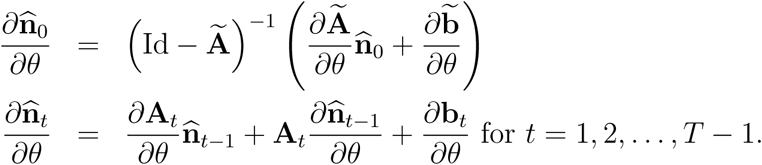

#### Uncorrelated fluctuations around a periodic signal

Stochastic variation in seasonal fluctuations or decadal oscillations can be modeled by combining the elements from the previous two examples. For descriptive purposes, assume there are *T* “seasons” per year and *t* counts the number of seasons that have elapsed. For each season 0 ≤ *i* ≤ *T* – 1, let **A**_*i*,1_, **A**_*i*,2_, **A**_*i*,3_, … and **b**_i,1_, **b**_*i*,2_, **b**_*i*,3_, … be i.i.d. sequences. For any time *t*, let *y*(*t*) = ⌊*t*/*T*⌋ correspond to the “year” and *s*(*t*) = *t* – *y*(*t*)*T* the “season.” Then we can write down a model of the following type:

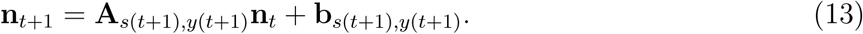

The periodic “deterministic skeleton” of this model is given by **n**_*t*+1_ = **Ā**_*s*(*t*+1)_**n**_*t*_ + **b̄**_*s*(*t*+1)_ where **Ā**_*i*_ = 𝔼[**A**_*i*, *t*_] and **b̄**_*i*_ = 𝔼[**b**_*i*, *t*_].

To understand the first two moments of the stationary distribution, we can look at the “yearly stroboscope” version of the model:

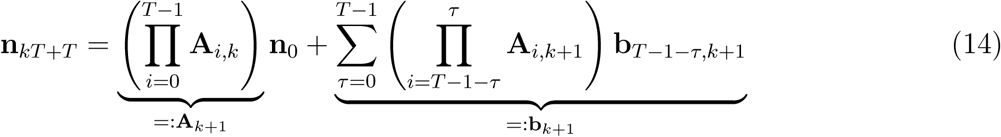

As **A**_1_, **A**_2_, … and **b**_1_, **b**_2_, … as defined above are i.i.d. sequences, the results for the i.i.d. environments can be applied here.

## Applications

To illustrate the applicability of the general methods, we apply them to a model of an unstructured population, a matrix model of reef corals [Hughes, 1984, Pascual and Caswell, 1991] and an integral projection model of giant clams [Yau et al., 2014].

### An unstructured, open population

The simplest stochastic, open model is for an unstructured population and takes the form

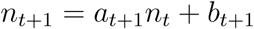

where *n_t_*, *a_t_*, and *b_t_* are scalars. A continuous-time version of this model, for example, was used by Gonzalez and Holt [2002] to describe open, sink populations in a fluctuating environments. We assume that *a_t_* > 0 and *b_t_* > 0 are independent, identically distributed sequences. More specifically, log *a_t_* and log *b_t_* are normally distributed with means *μ_a_*, *μ_b_*, variances 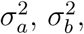 and correlation *ρ*.

This model has a unique stationary distribution provided that *r* = 𝔼[log *a_t_*] = *μ_a_* < 0 (Fig. 1). The mean of this stationary distribution is finite if the arithmetic mean 𝔼[*a_t_*] of *a_t_* is less than one. Namely,

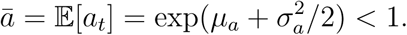

**Figure 1:**
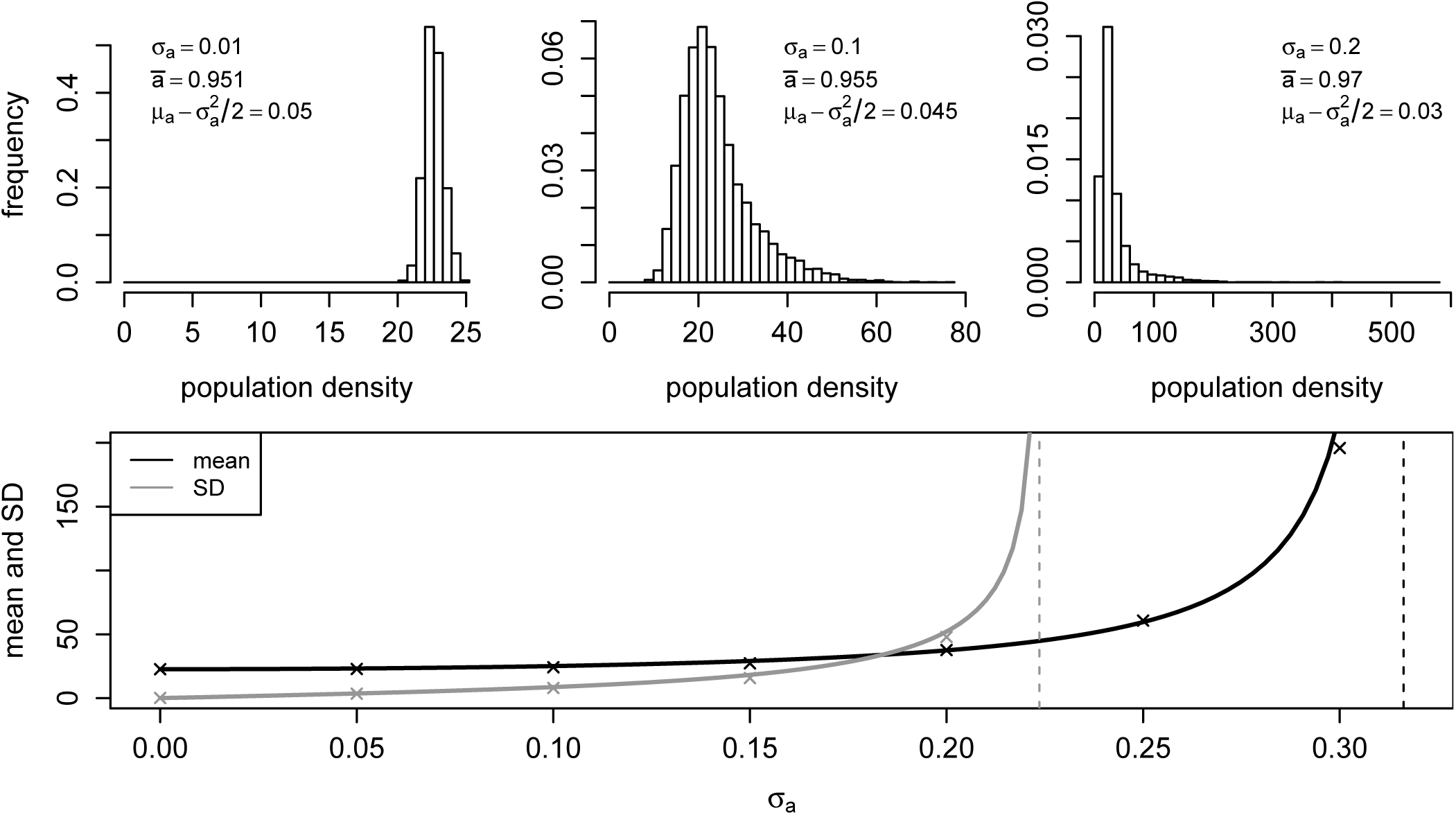
The effect of increasing within patch fluctuations *σ_a_* on the mean and stationary distribution of the scalar model *n*_*t*+1_ = *a*_*t*+1_*n_t_* + *b*_*t*+1_. In top panels, numerical approximations of stationary distributions for three levels of local fluctuations *σ_a_*. In the bottom panel, the mean and standard deviation of the stationary distribution as a function of *σ_a_*. Solid line corresponds to the analytic formulas (15)–(16), and the crosses correspond to mean and standard deviation of the numerically approximated stationary distribution. For the numerical approximations, the process was simulated for 100, 000 years. Parameters: *a_t_* is log-normally distributed with log mean *μ_a_* = − 0.05 and standard deviation *σ_a_* as shown. *b_t_* is log-normally distributed with log mean *μ_b_* = 0.05 and log standard deviation *σ_b_* = 0.05. *a_t_* and *b_t_* are independent.

Equivalently, 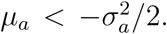 As shown in the top panels of Figure 1, as 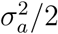 approaches *μ_a_*, the stationary distribution becomes highly skewed with a “thick tail.” When it is well-defined, the mean of stationary distribution equals

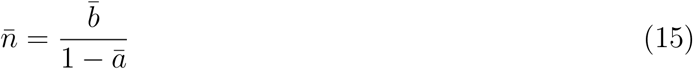

where *b̄* = 𝔼[*b_t_*] = exp(*μ_b_* + 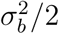). This stationary distribution has a finite variance if

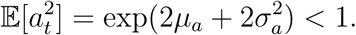

Equivalently, *μ_a_* < 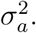 Hence, **n̂** has a finite mean but infinite variance whenever – 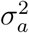 < *μ_a_* < –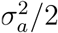 (Fig. 1). When the variance is finite, it equals

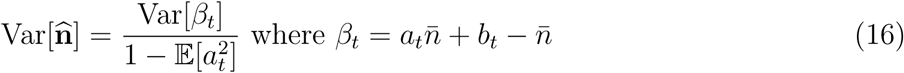

and

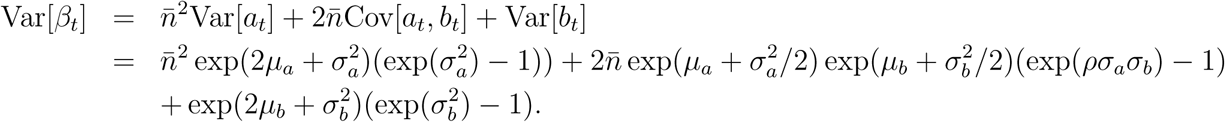

As 𝔼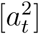 = *ā*^2^ + Var[*a_t_*], this expression for Var[**n̂**] implies that variation in the local dynamics (Var[*a_t_***n̂**]) contributes more to the variation in the population densities (Var[**n̂**]) than the variation in external recruitment (Var[*b_t_*]) i.e. Var[*a_t_*] and Var[*b_t_*] appear in a symmetric manner in the numerator but the denominator decreases with Var[*a_t_*]. Moreover, positive correlations between local dynamics and external recruitment increase the variance in the long-term population densities.

### Matrix model of reef corals with space-limited recruitment

We continue with a case study of an open population of reef corals with space-limited recruitment. Corals are sessile organisms with pelagic larvae. As such, recruitment isn’t strongly affected by local coral populations, but rather upon reproductive subsidies from external populations. Hughes [1984] first developed a size-structured, matrix population model of corals in a closed population. This model was extended by Pascual and Caswell [1991] to allow for open populations and space-limited recruitment. The model of Pascual and Caswell [1991] drew heavily upon the theoretical framework developed by Roughgarden et al. [1985] to describe open, age-structured barnacle populations with space-limited recruitment. We extend the model of Pascual and Caswell [1991] here to include stochastic recruitment.

Let **n**_*t*_(*i*) be the number of organisms is size class *i* at time *t*. At each time step, surviving individuals either remain in their current size class with probability *R*(*i*), or grow to the next size class with probability *P*(*i*). The average size, measured as an area, of each organism in a given size class is given by *a*(*i*). New individuals enter the population at a rate proportional to the amount of free space available, where the settling parameter, *s*_*t*+1_, is the number of larvae per unit area that enter the population and survive until the next census. If *α* is the total area of suitable substrate, then the amount of free, unoccupied substrate at each time step given by

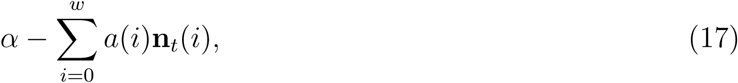

where *w* is the total number of size classes. The full model is then given by

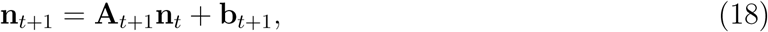

with

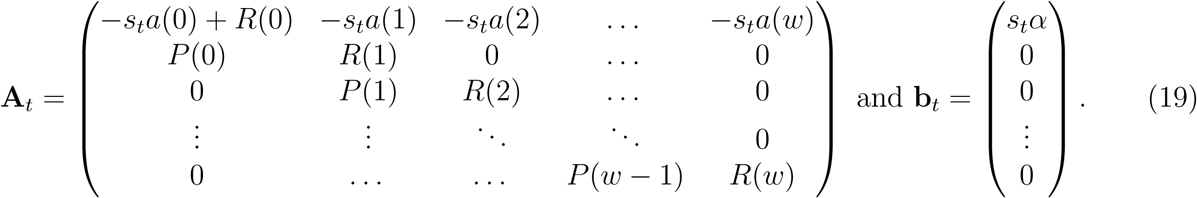

To explore this model, we use the parameter values of Pascual and Caswell [1991] (adapted from Hughes [1984]). These parameter values were obtained from data for the reef coral *Agaricia agaricites* from 1978 to 1979. Corals were groups into three size classes: 0-10 cm^2^, 10-50 cm^2^, and 50-200 cm^2^. The total amount of available substrate is set equal to 12 m^2^. With these choices, the matrix model becomes

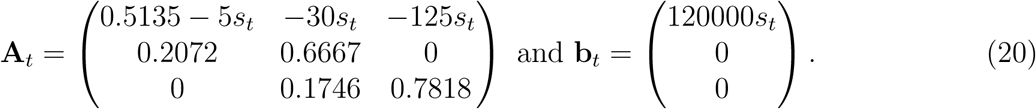

We assume that settlement rates vary randomly between a low settlement rate of *s_ℓ_* = 0.01 and a high settlement rate of *s_h_* = 0.04. These values lie below the critical settlement value of *s* = 0.06, above which the increased settlement rate leads to unstable and unbounded dynamics [Pascual and Caswell, 1991]. Let *p* be the frequency of a high settlement year, with low settlement years occurring with frequency 1 – *p*.

When the frequency of good years is low (*p* = 0.05), the stationary distribution is concentrated near the stable size distribution for the deterministic model with only bad years (*p* = 0; top panels in Fig. 2). At intermediate frequencies of good years (*p* = 0.5), the stationary distribution becomes increasingly normally distributed from the smallest to largest size classes (bottom panels in Fig. 2). The population exhibits the greatest mean densities when the frequency of good years is high (Fig. 3A), and the greatest variation in abundances in all size classes when good years occur at an intermediate frequency (*p* ≈ 0.4 in Fig. 3B). Intuitively, at high or low frequencies of good years, the dynamics are nearly deterministic and, consequently, exhibit minimal variation. Adjacent size classes exhibit weak, positive correlations in densities, while the largest and smallest size classes exhibit negative correlations (Fig. 3C). As higher frequencies of good years leads to lower availability of substrate, these correlations are less positive or more negative at higher frequencies of good years. One year autocorrelations are greatest for the largest size class, and least for the smallest size class (Fig. 3D). Intuitively, stochastic recruitment only occurs in the smallest size class which reduces the correlation with densities in earlier years. Increasing frequency of good years make these stochastic recruitment events more consistent and, thereby, increase autocorrelations in the smallest size class.

**Figure 2:**
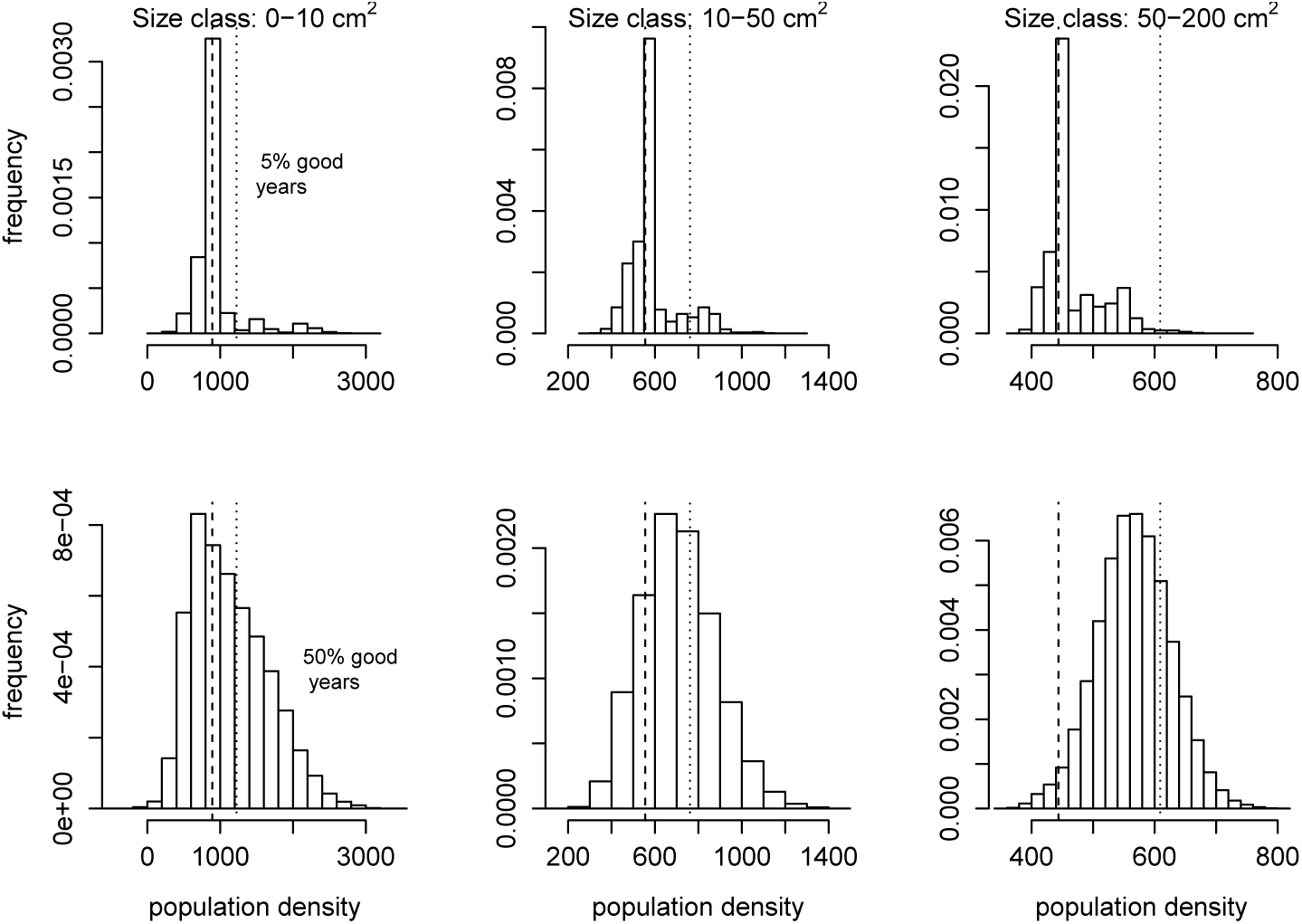
Numerical approximations of the stationary distribution, using a single long run of 100, 000 years, for low frequencies (*p* = 0.05) and intermediate frequencies (*p* = 0.5) of good years. Dashed vertical lines indicate the population density for each size class with 0% good years (*p* = 0), while dotted vertical lines indicate the population density for each size class with 100% good years (*p* = 1). Parameters as described in the main text.

**Figure 3:**
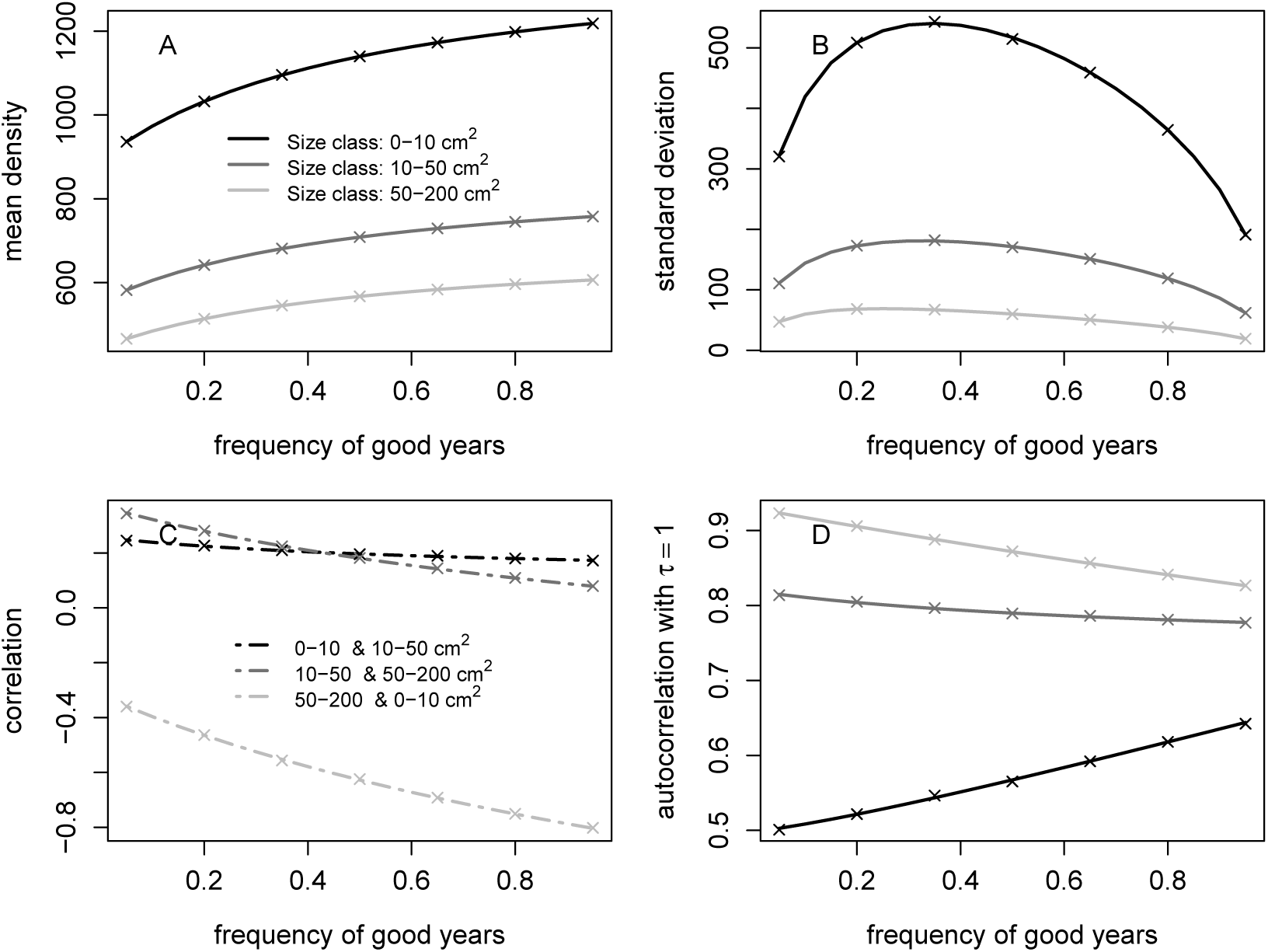
The effect of frequency of good years on mean density, standard deviations within size classes, correlations between size classes, and temporal autocorrelation within size classes. Solid lines come from the analytic formulas and × correspond to simulation estimates. Numerical simulations were run for 100, 000 years. Parameters as described in the main text.

To illustrate the use of the sensitivity and elasticity formulas, we examine elasticities of the mean and standard deviation of density with respect to the densities of recruits arriving in good and bad years i.e. *s_h_* and *s_ℓ_*. The details for computing the sensitivities and elasticities are presented in Appendix S5. The mean and standard deviation in density is most sensitive to the density of recruits arriving in bad years, *s_ℓ_*, particularly when the frequency of good years is low (Fig. 4). Interestingly, the elasticity of mean densities is not stage-dependent. The elasticity of the standard deviation is negative with respect to the density of recruits arriving in low recruitment years. The elasticity of the standard deviation with respect to the density of recruits arriving in high recruitment years does depend slightly on stage, with smaller size classes showing a higher increase in standard deviation with changes in *s_h_*.

**Figure 4:**
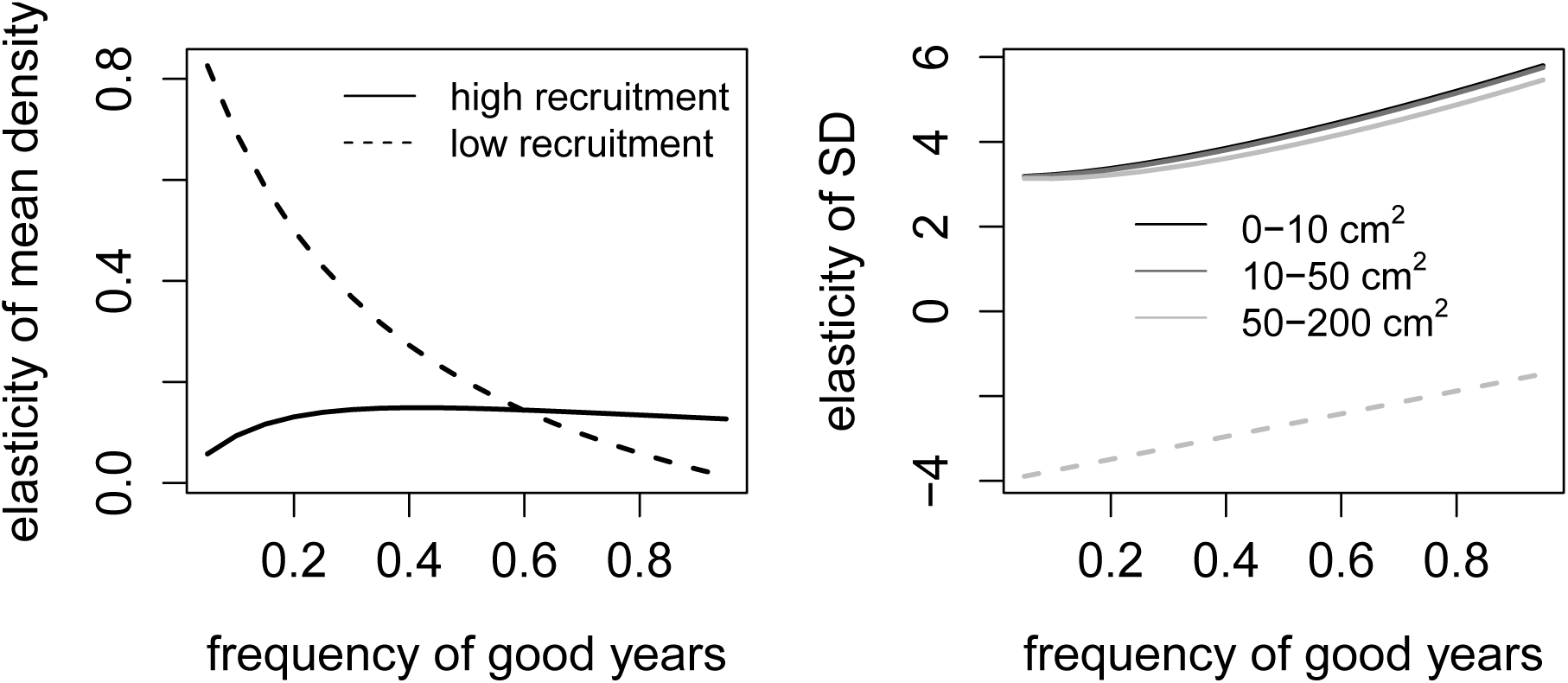
The effect of the frequency of good years on the elasticity of the mean and standard deviation of coral densities with respect to the densities of recruits arriving in a good (solid line) and bad (dotted line) year. Parameters as described in the main text.

### Integral projection model of giant clam dynamics

In this example, we apply the above methods to a population of giant clams, *Tridacna maxima*, using data from Yau et al. [2014]. Yau et al. [2014] developed two deterministic size-structured IPMs to describe a population of giant clams on Mo’orea, French Polynesia. One IPM assumed a completely closed population with no external recruitment, while the second IPM assumed a completely open population with no local retention. Yau et al. [2014] also investigated the amount of local retention that was required for the population to persist in the absence of external recruitment.

To parameterize the model, Yau et al. [2014] measured demographic rates of 1,949 clams annually from 2006 to 2010. Survival data was fit using nonlinear logistic regression, while growth data was fit using ordinary least squares regression of size at time *t* + 1 on size at time *t*. The variance in growth was also allowed to vary as a function of size. The size of new recruits was estimated using data on the size of clams < 50 mm counted each year. Recruit size was assumed to be normally distributed with mean and standard deviation estimated from the data. For the open model, the number of recruits was set equal to the average number of clams < 50 mm that were counted each year. For the closed model, the number of recruits was assumed to be proportional to the size-specific adult gonadal mass, with only individuals > 66.1 mm contributing reproductively to the population. Gonadal mass was converted to new recruits using a conversion factor, *c_f_*, which was estimated by averaging the observed ratio of recruits to total gonadal mass each year. Finally, to determine the amount of local retention required for persistence, Yau et al. [2014] varied *c_f_* to find the threshold such that the population transitioned from self-sustaining (population growth rate λ > 1) to declining (λ < 1). Table 1 shows the demographic functions and parameters used in the IPM.

**Table 1:**
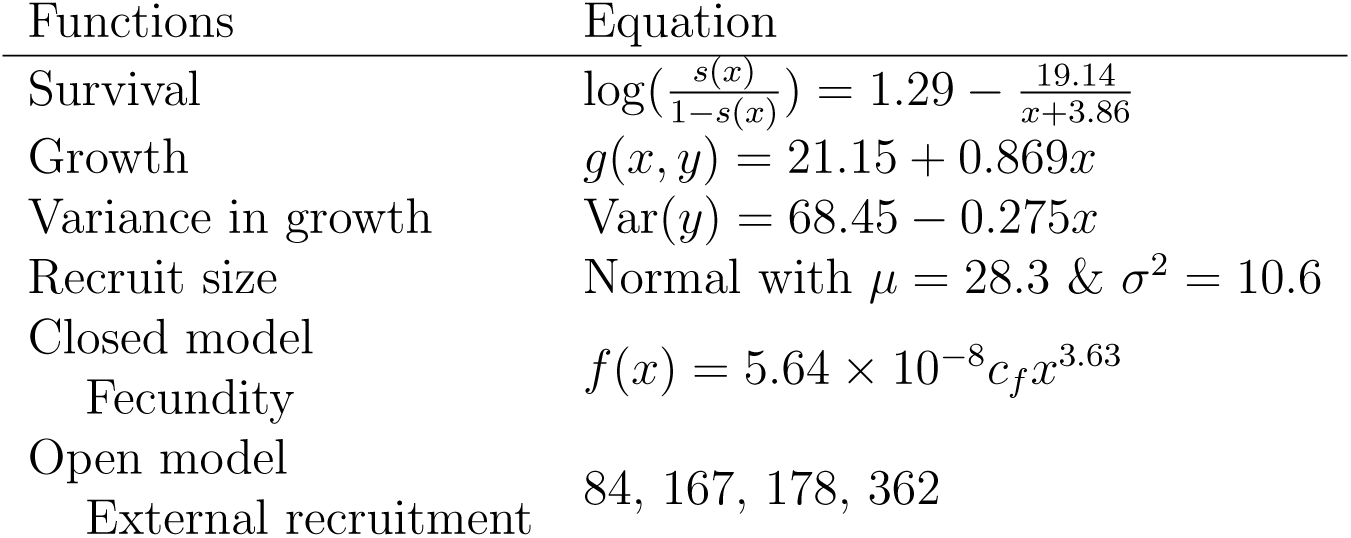
Demographic functions and parameters used in the IPM, from Yau et al. [2014]. The variable *x* represents the size of a clam, in mm. The parameter *c_f_* is a conversion factor that converts grams of dry gonad mass to new recruits, and provides an indication of the degree of local retention, with *c_f_* = 0 indicating no local retention.

For the stochastic IPM, we assume a population that includes both local retention and external recruitment. The model is written as

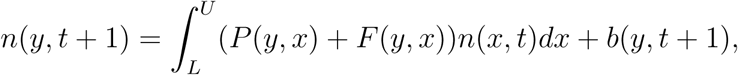

where *P*(*y*, *x*) represents the survival and growth of individuals from size *x* to size *y*, *F*(*y*, *x*) represents the local recruitment, and *b*(*y*, *t* + 1) represents the external recruitment. Clam sizes range from *L* = 1 to *U* = 200 mm. To incorporate stochasticity into the model, we generated four external recruitment vectors, one for each year of collected data: *∫ b*_1_(*x*) *dx* = 84, *∫ b*_2_(*x*) *dx* = 167, *∫ b*_3_(*x*) *dx* = 178, and *∫ b*_4_(*x*) *dx* = 362. Each *b* vector is assumed to occur with equal probability. Additionally, we varied the degree of local retention, *c_f_* from *c_f_* = 0 (a fully open population with no local retention) to *c_f_* = 0.78 (a population with λ just below 1 in the absence of external recruitment).

Figure 5 shows the size-specific mean density, coefficient of variation, and correlation between size at time *t* and size at time *t* +1 for *c_f_* = 0 (top panel) and *c_f_* = 0.7389 (middle panel). While the size-distribution remains roughly the same for different levels of *c_f_*, overall more clams are present when local retention is higher. As expected, the coefficient of variation is highest for the smallest sized individuals that receive the greatest amount of stochastic external recruitment. Additionally, as *c_f_* increases and the deterministic portion of the model becomes greater, the correlation across all sizes also increases, as expected.

**Figure 5:**
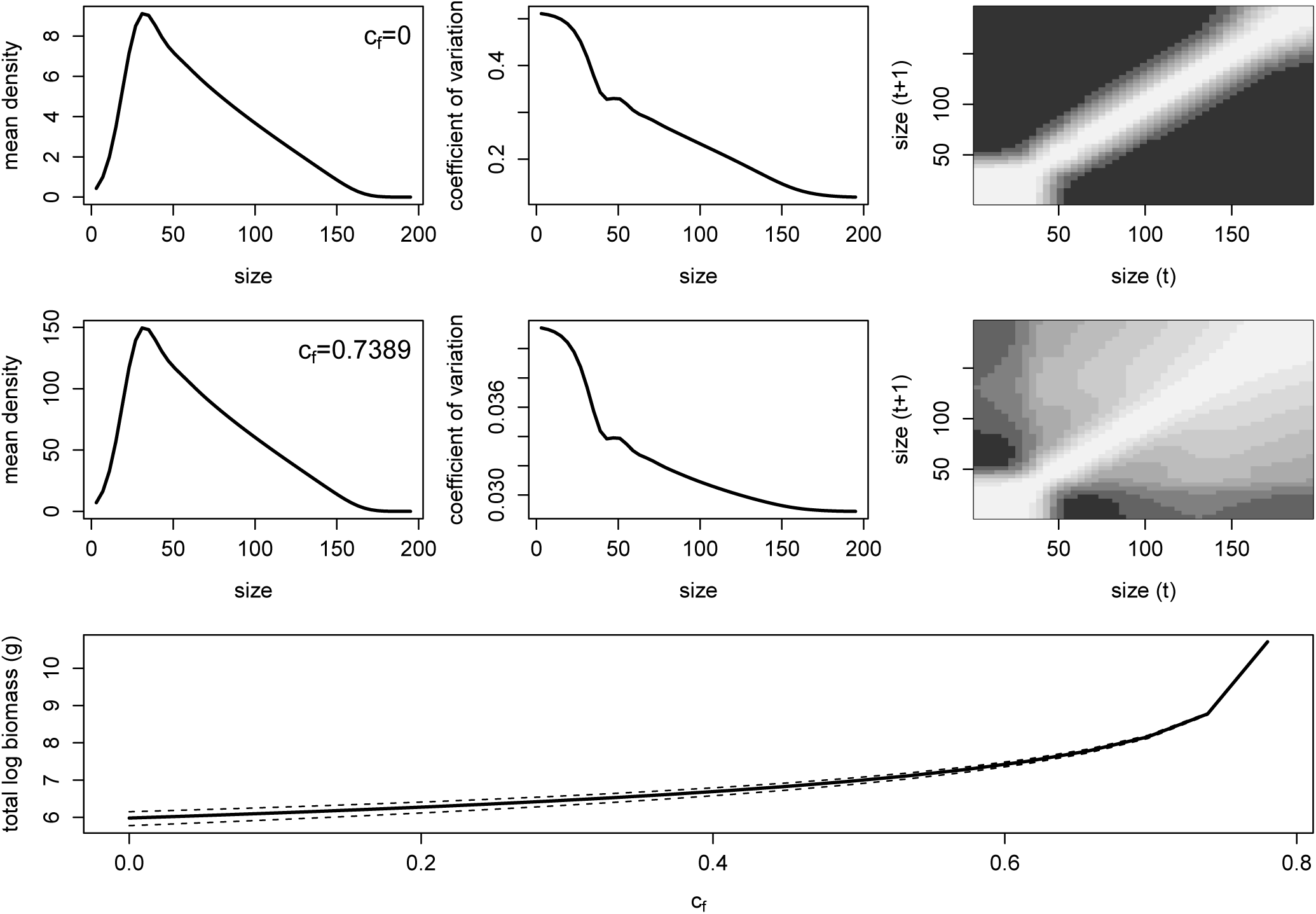
The size-specific mean density, coefficient of variation, and correlation between size at time *t* and size at time *t* + 1 in an open population with no local retention (*c_f_* = 0; top panel), and a clam population with λ just below 1 in the absence of external recruitment (*c_f_* = 0.7389; middle panel). The bottom panel shows the total log clam biomass as a function of the degree of local retention, *c_f_*.

We also investigated how the expected clam biomass, *B*, changes as a function of local retention. The expected total biomass of the population is equal to 𝔼[*B*] = 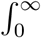 *w*(*x*)𝔼[**n̂**(*x*)], where *w*(*x*) is a scaling parameter that converts an individual of size *x* to its biomass. The variance of the biomass is equal to var[*B*] = *∫∫ w*(*x*)*w*(*y*)**Cov**[**n̂**(*x*), **n̂**(*y*)]*dxdy*. The bottom panel of Fig. 5 shows the log of the expected biomass across the range of *c_f_* values, as well as the log of the standard deviation. As expected, as the degree of local retention increases such that the closed population growth rate λ nears 1, the log biomass increases substantially. Additionally, the variance in biomass decreases as *c_f_* increases.

We examined the sensitivity of the mean density and the standard deviation of the mean density to the probability of a particular recruitment year (Fig. 6). We assume that if the probability of type *i* year, *p_i_*, is increased by *θ*, then the probability of the remaining three year types is decreased by *θ*/3. The details for computing these sensitivities are presented in Appendix S5. Our computations reveal that the mean density for all sizes is most sensitive to the frequency of the highest recruitment years and slightly less sensitive to the frequency of the lowest recruitment years (darkest versus lightest curves in the left panel of Fig. 6). Mean densities are much less sensitive to the frequency of the intermediate recruitment years. In contrast, the standard deviation of the densities in all sizes are least sensitive to the frequency of the lowest recruitment years and much more sensitive to frequencies of all the other types of recruitment years (lightest curve versus all other curves in the right hand panel of Fig. 6). In all cases, the sensitivities follow the size distribution of the population.

**Figure 6:**
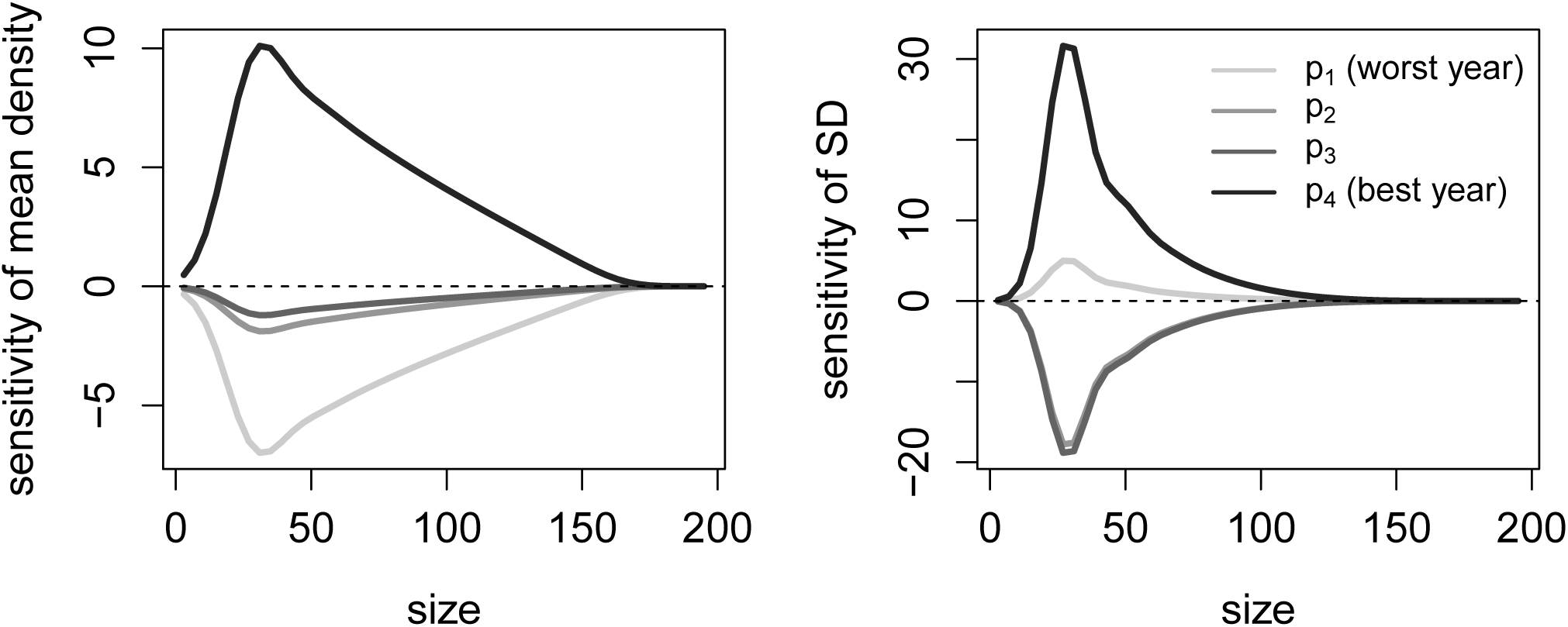
The sensitivity of the mean density and the standard deviation of the mean density to changes in the frequency of a particular year, *p_i_*, with *p*_1_ representing the frequency of the worst year, and *p*_4_ representing the frequency of the best year. We assume that if the probability of type *i* year is increased by *θ*, then the probability of the remaining three year types is decreased by *θ/*3.

## Discussion

Here, we develop methods for analyzing affine models of open populations with continuous and discrete population structure (e.g. size and stage) experiencing stochastic local demography and stochastic external recruitment. Provided that the local demography is unable to support the population (i.e. the local asymptotic growth rate is negative due to mortality and emigration exceeding reproduction), we show that all initial population states converge to the same asymptotic behavior. When the environmental fluctuations are non-stationary, this limiting behavior corresponds to a “pull-back attractor” in the mathematical theory of non-autonomous systems [Kloeden and Rasmussen, 2011] which more recently has been dubbed an “asymptotically environmentally dependent trajectory” by Chesson [2017]. When the environmental fluctuations are stationary, these limiting behaviors are characterized by a unique stationary distribution of the model. For environments with a mixture of periodic and serially uncorrelated fluctuations, we derived explicit formulas for the mean and covariance of these stationary distributions and provide sensitivity sensitivity formulas for these quantities. In particular, our results extend the work of [Gross and Edmunds, 2015] for discretely structured populations experiencing uncorrelated environments in only external recruitment.

We show, using a general, unstructured model, that variation in local dynamics contributes more to variance in population densities relative to variance in external recruitment. We show also that positive correlations between local dynamics and external recruitment increase the variance in long-term population densities, while negative correlations will decrease this variance. Finally, we show that, under certain conditions, it is possible for a population to be regulated, but have an infinite mean density. This effect occurs when the stochastic growth rate of the closed portion of the model is negative, but the log eigenvalue of the averaged system is positive. In this case, large, rare fluctuations contribute to the mean, but since the stochastic growth rate of the closed portion of the model is negative, the population remains regulated. This result provides an open population parallel to what Lewontin and Cohen [1969] found for closed populations: “even though the expectation of the population size may grow infinitely large with time, the probability of extinction may approach unity, owing to the difference between the geometric and arithmetic mean growth rates.”

We also show the application of the methods to two structured marine populations, specifically investigating the impact of changing the frequency of good recruitment years, and changing the degree of openness of the population. In general, our results are intuitive. Mean population densities are highest when there is less variability in recruitment, while the standard deviation of the mean population density is highest when there is greater variability in recruitment. With the IPM, we find that increases in mean density only occur with increasing the frequency of the best recruitment year, and that this effect is slightly greater than the negative effect of increasing the frequency of the worst recruitment year, likely due to relative distance of the best and worst recruitment levels from the average recruitment value. Moreover, using the matrix model, we find that a small number of good years is nearly as effective as a large number of good years at contributing to increases in mean densities, due to the dependence of recruitment on available open space.

Of particular interest is the use of the methods to study the covariance structure of the population. Using the IPM, we see that decreasing recruit variability by decreasing the degree of population openness leads to higher correlations across sizes of individuals in the population. Conversely, when recruit variability is high, correlation across sizes is reduced, and is mostly through direct growth from one size to another between time steps. In the matrix model, correlation between sizes is positive for adjacent size classes (i.e. high population densities in one size class will lead to high densities in the next size class). However, the correlation between the smallest and largest size class is negative, with the relationship most negative when the frequency of good years is high. This is most likely due to the impact of space-limitation on recruitment. Finally, we can use the methods to investigate the autocorrelation structure of the population between different time steps. For a single-year time step in the matrix model, we see that the autocorrelation is highest for the largest size classes (with no variable input), and lowest for the smallest size class (due to the variability in recruitment). As the frequency of good years increases, the autocorrelation of the smallest size class increases substantially.

While we used the methods presented here for models of open populations, they also apply to two other classes of models. First, consider a nonlinear, stochastic difference equation **n**_*t*+1_ = *F*(**n**_*t*_, *ξ_t_*) where *ξ_t_* describe the environmental fluctuations. If these fluctuations are “small” and **n̂** is a stable equilibrium of the difference equation **n**_*t*+1_ = *F*(**n**_*t*_, 0), then the stochastic dynamics near this equilibrium may be approximated by **n**_*t*+1_ = *∂*_n_*F*(**n̂**, 0)(**n**_*t*_ – **n̂**) + *∂_ξ_F*(**n̂**, 0)*ξ_t_* which is a model of the form considered here. Hence, our methods provide a means to approximate the covariance matrix of this linear approximation whenever the *ξ_t_* are serially uncorrelated. Second, our methods also apply to extensions of multispecies models studied by [Ives et al., 2003, Cooper et al., 2015, Gross and Edmunds, 2015]. If *x_i_* denotes the density of species *i*, these models take the form *x_i_*(*t* + 1) = *r_i_*(*t*)*x*_1_(*t*)^*a*_*i*l_(*t*)^*x*_2_(*t*)^*a*_*i*2_(*t*)^… *x_n_*(*t*)^*a*_*ik*_(*t*)^ where *r_i_*(*t*) corresponds to intrinsic growth rates of species *i* and *a_ij_* (*t*) describes time-dependent species interactions. Setting *n_i_* = log *x_i_* yields a model of the form **n**_*t*+1_ = **A**_*t*_**n**_*t*_ + **b**_*t*_ where **A**_*t*_ = *a_ij_*(*t*) and **b**_*t*_ = log *r_i_*(*t*). Hence, our results apply and generalize the work of [Ives et al., 2003, Cooper et al., 2015, Gross and Edmunds, 2015] who assumed that the interaction terms did not vary in time.

In conclusion, when studying real-world populations, it’s important to understand the interplay of open recruitment, stochasticity in demographic rates, and population structure. This is particularly important when studying organisms with ecological, economic, or public interest importance, such as invasive or threatened species, and when studying the effect of changing environmental conditions (for instance, due to climate change). Our results show that, for open populations, it is important to understand the effects of local demographic stochasticity, rather than focusing solely on variability in recruitment, as is often done, particularly for marine organisms. We also show that our methods can be used to directly assess the impact of decreases in the frequency of good years, as well as investigate the correlation between population states, and changes in population states due to changing environmental conditions.

In these Appendices we present the proofs of convergence for the non-autonomous case, existence and convergence to a stationary distribution for stationary environments, the derivation of mean and covariance of the stationary distribution for uncorrelated environments, and identification of the periodic solution for periodic environments. For the uncorrelated and periodic environments, we derive the sensitivity formulas presented in the main text.

## Appendix S1 Convergence in general

In this Appendix, we consider solutions of (1) with minimal assumptions on the sequence **A**_1_, **A**_2_, … and **b**_1_, **b**_2_, …. In particular, the sequence need not be stationary. To prove exponential convergence of all initial conditions to the same time varying solution, we assume that

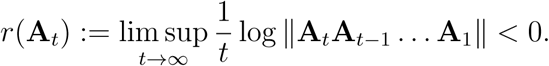

Given two initial conditions **n**_0_ and **ñ**_0_, let **n**_*t*_ and **ñ**_*t*_ be the corresponding solutions. Then

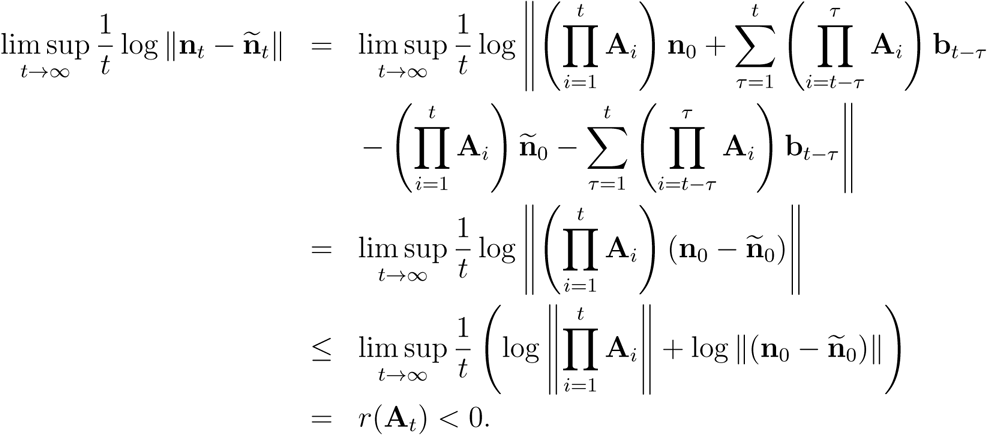

Thus, as claimed, solutions corresponding to different initial conditions converge to one another at an exponential rate.

## Appendix S2 Convergence for stationary environments

Assume that **A**_1_, **A**_2_, **A**_3_, … are a stationary and ergodic sequence of non-negative operators, and 𝔼[max{log ∥**A**_*t*_∥, 0}] < ∞. As log ∥**A**_*t*+*s*_ … **A**_1_∥ ≤ log ∥**A**_*t*+*s*_ … **A**_*s*+1_∥ + log ∥**A**_*s*_ … **A**_1_∥, Kingman [1973]’s subadditive ergodic theorem implies there exists an *r* (possibly − ∞) such that

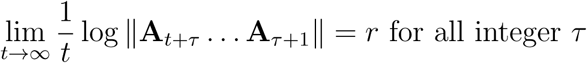

with probability one. Let …, **A**_−2_, **A**_−1_, **A**_0_, **A**_1_, **A**_2_, … and …, **b**_−2_, **b**_−1_, **b**_0_, **b**_1_, **b**_2_, … be the unique extensions to bi-infinite, stationary, ergodic sequences. Kingman’s subadditive ergodic theorem also implies

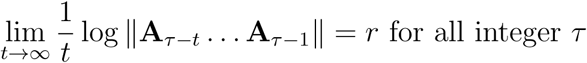

with probability one.

Now assume that 𝔼[log ∥*b_t_*∥] < ∞ and *r* < 0. Define

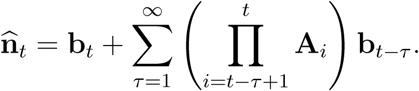

We will show that **n**_*t*_ is well-defined with probability one, stationary, and **n**_*t*_ converges to **n̂**_*t*_ any initial condition **n**_0_. Our arguments follow Brandt [1986] who proved these statements for the scalar case (i.e. **n**_*t*_ takes values in [0, ∞)).

To see that **n̂**_*t*_ is will defined with probability one, notice that

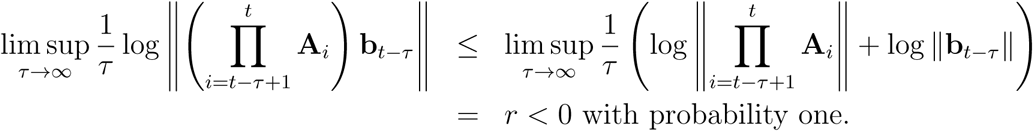

Hence, Cauchy’s criterion for convergence of infinite sums, **n̂**_*t*_ is well-defined with probability one.

Next we verify that **n̂**_*t*_ is a solution to the stochastic difference equation:

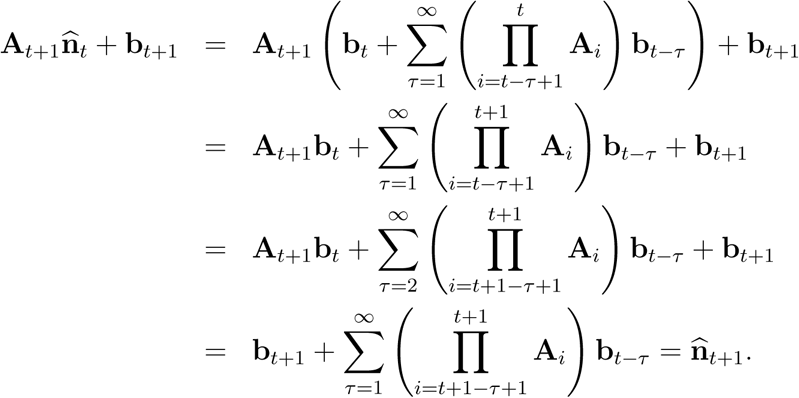

Stationarity of **n̂**_*t*_ follows from stationarity of **A**_*t*_ and **b**_*t*_.

To prove convergence of **n**_*t*_ for a given initial state **n**_*t*_ for a given initial state **n**_0_ to **n̂**_*t*_ as *t* → ∞, notice that

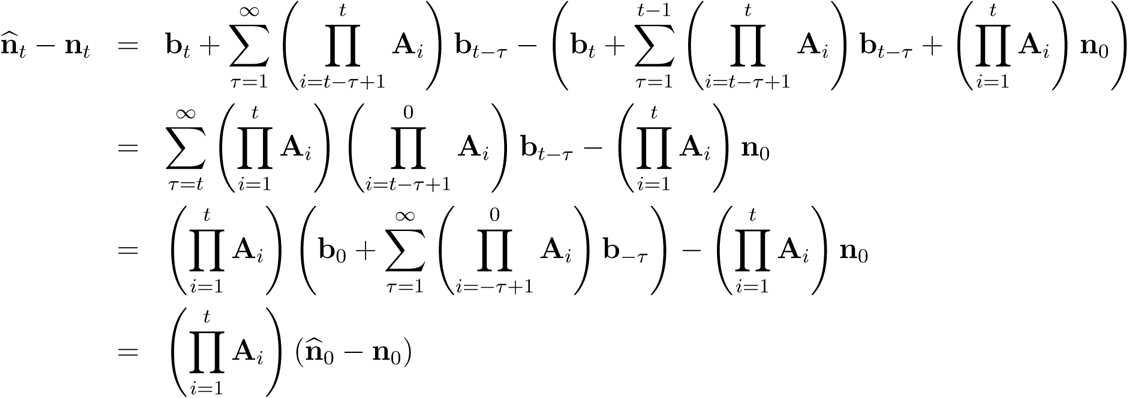

Hence,

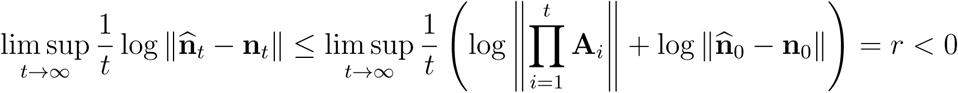

with probability one. Hence, as claimed, **n**_*t*_ converges exponentially fast to **n̂**_*t*_.

## Appendix S3 Formulas for uncorrelated environments

Assume that **A**_1_, **A**_2_, … and **b**_1_, **b**_2_, … are independent and identically distributed sequences. Furthermore, assume 𝔼[log max{∥**b**_*t*_∥, 0}] < ∞ and *r* < 0. Then Appendix S2 implies there exists a stationary solution **n̂**_*t*_.

To find the expectation of **n̂**_*t*_ (under the assumption it exists), define

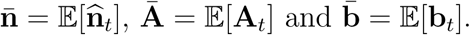

By stationarity and independence, we have that

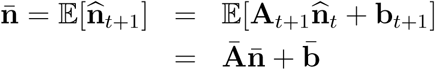

Thus,

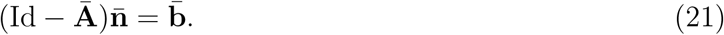

If the spectral radius of **Ā** is less than one, then Id − **Ā** is invertible and

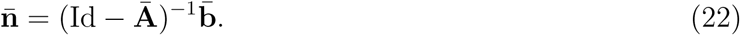

To determine the sensitivity of **n̄** to a parameter *θ*, implicit differentiation of (21) yields

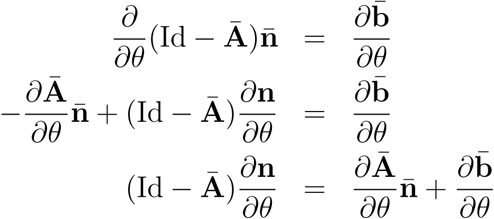

Thus, provided the spectral radius of **Ā** is less than one,

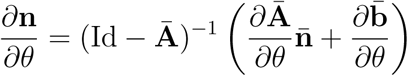

as claimed in the main text.

To find the covariance of **n̂**_*t*_ (assuming it exists), define

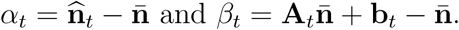

Then 𝔼[*β_t_*] = 0, 𝔼[*α_t_*] = 0, and **Cov**[**n̂**] = 𝔼[*α_t_* ⊗ *α_t_*]. Stationarity, independence, and properties of tensor products imply

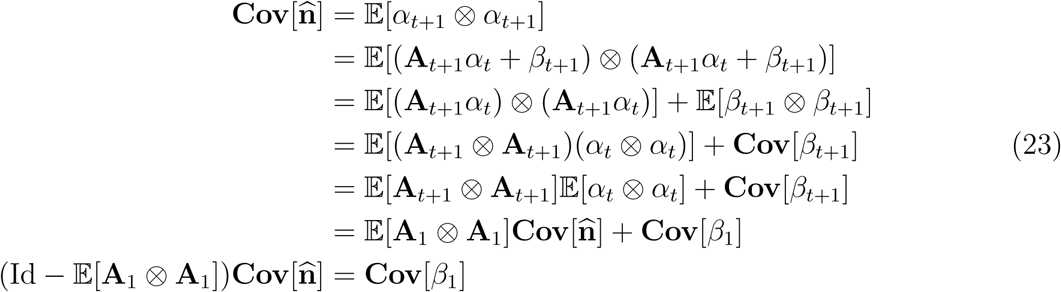

Hence, provide that the spectral radius of 𝔼[**A**_1_ ⊗ **A**_1_] is less than one, then

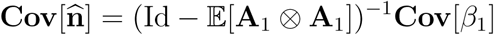

as claimed in the main text. In the finite dimensional case,
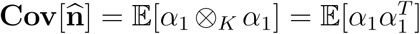
where ^*T*^ denotes the transpose. We can solve this system of linear equations using the vec operator and the Kronecker product as follows:

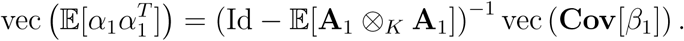

To find the sensitivity formula for **Cov**[**n̂**], we can implicitly differentiate (23) with respect to the parameter *θ*:

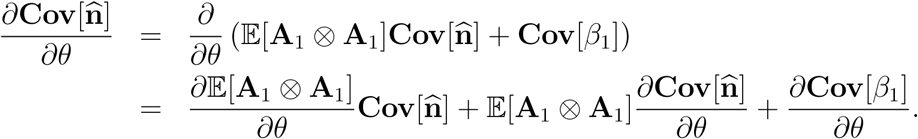

Solving yields

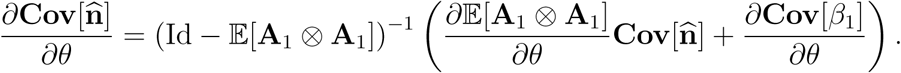

In finite dimensions, this equation can be expressed in terms of the vec and Kronecker operations as presented in the main text.

Finally, the covariance between **n̂**_0_ and **n̂**_*t*_ with *t* ≥ 1 is given by

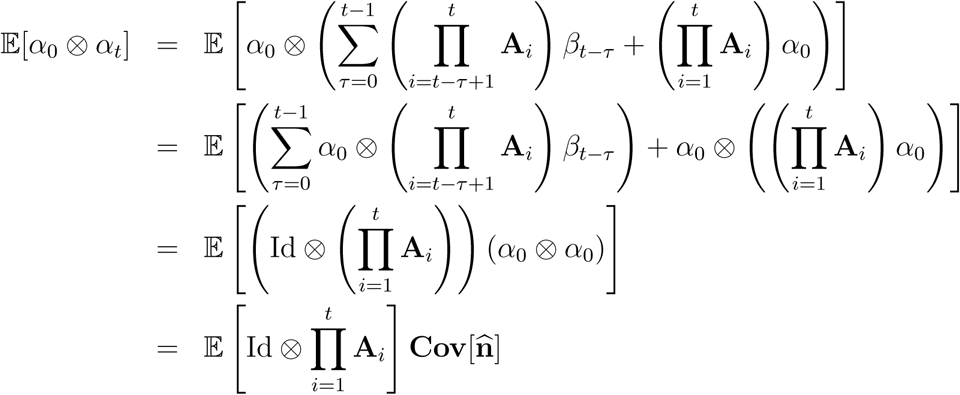

as claimed in the main text. The sensitivity formula in the main text follows from direct differentiation using the product and chain rules.

## Appendix S4 Formulas for periodic environments

To find the formula for this periodic trajectory, we consider the time *T* map given by

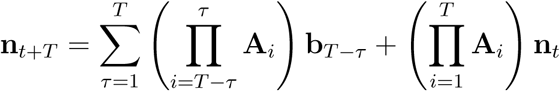

whose equilibrium **n̂** satisfies

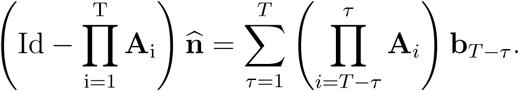

As the dominant eigenvalue of
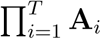
is less than one, the operator
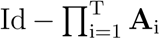
is invertible and we have

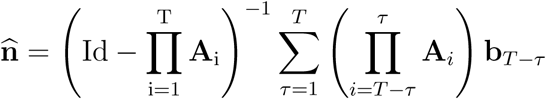

If we define **n̂**_0_ = **n̂** and **n̂**_*t*_ = **A**_*t*_**n̂**_*t*−1_ + **b**_*t*_ for 1 ≤ *t* ≤ *T* − 1, then for any initial condition **n**_0_ and 0 ≤ *τ* ≤ *T* − 1, we have

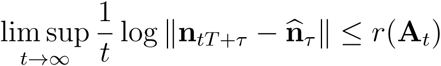

i.e. all solutions converge exponentially quickly to the periodic solution **n̂**_0_, …, **n̂**_*T*−1_.

## Appendix S5 Calculations for examples

For the coral reef and giant clam examples, the **A**_*t*_ and **b**_*t*_ are chosen at random from a finite number of choices, say **A**(1), …, **A**(*ℓ*) and **b**(1), …, **b**(*ℓ*), with probabilities *p*(1), …, *p*(*ℓ*). For these types of models,

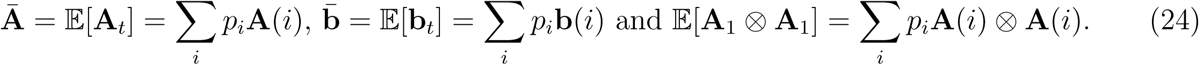

For the sensitivity formulas, there are two cases to consider. First, suppose that the **A**(*i*) and **b**(*i*) depend smoothly on some parameter *θ*, but *p_i_* do not depend on *θ*. Then

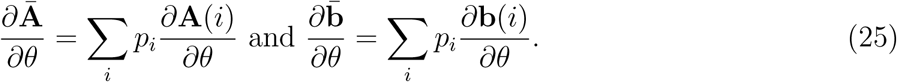

Furthermore,

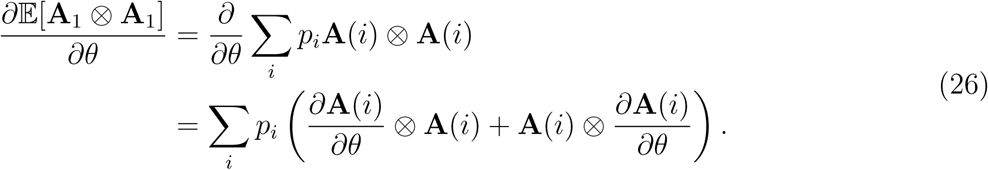

and

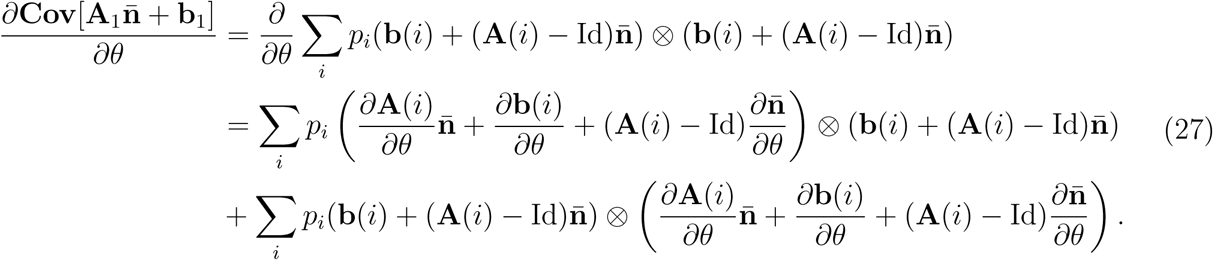

Second, suppose that the *p_i_* depend smoothly on the parameter *θ*, but **A**(*i*) and **b**(*i*) do not depend on *θ*. Then

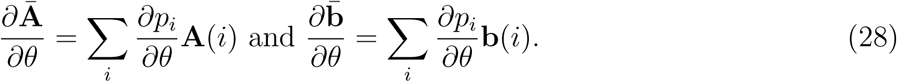

Furthermore

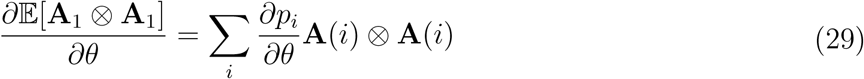

and

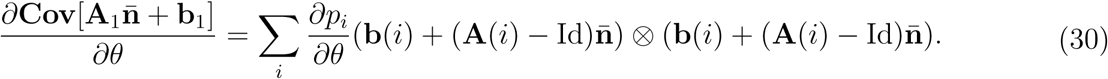

For the coral reef example, let *p*_1_ = *p*, *p*_2_ = 1 − *p*,

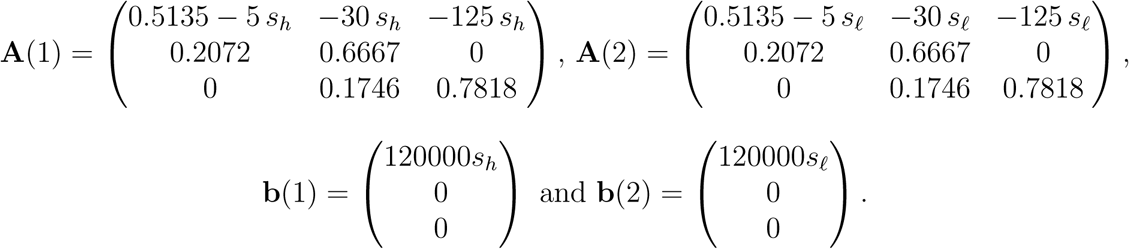

Thus,

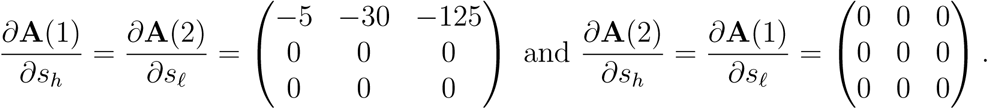

Moreover,

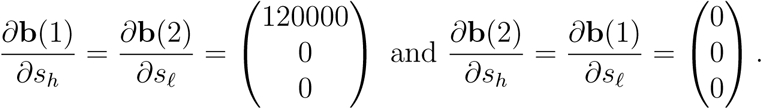

With these derivatives and formulas (24)–(27), we can use the sensitivity formulas in the main text to compute the sensitivities of **n̄**, **Cov**[**n̂**], and **Cov**_*τ*_ [**n̂**] for the coral reef model to parameters *θ* = *s_h_* and *θ* = *s_ℓ_*.

For the giant clam IPM example, we have *p*1, …, *p*4, **A**(1) = ⋯ = **A**(4) = **A**, and **b**(1), …, **b**(4). We calculated the sensitivities of means and covariances to the frequencies of different recruitment values. Specifically, to model an increase in the frequency of type *i* years, we let replaced *p_i_* by *p_i_* + *θ* and *p_j_* for *j* ≠ *i* by *p_j_* − *θ/*3. Then

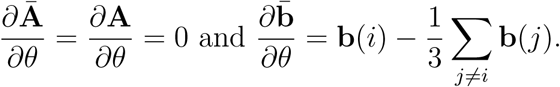

Furthermore,

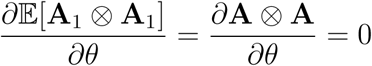

and

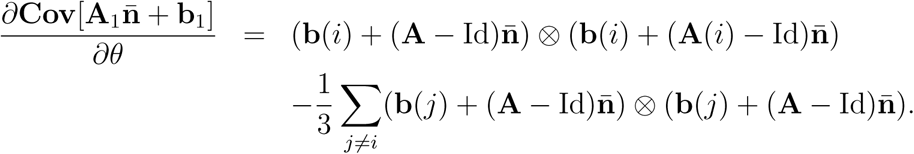

